# Non-Pyrogenicity and Biocompatibility of Parylene-Coated Magnetic Bead Implants

**DOI:** 10.1101/2023.09.01.555995

**Authors:** Cameron R. Taylor, Joshua K. Nott, Nilmini H. Ratnasena, Joel M. Cohen, Hugh M. Herr

## Abstract

We verify the non-pyrogenicity of magnetic bead implants and submit them to a full chemical characterization and toxicological risk assessment. Further, we validate the cleaning efficacy, steam sterilization, and dry time of a magnetic bead insertion device. We believe these results to be valuable to further scientific progress in the use of magnetic bead implants for human-machine interfacing.

## Introduction

In the manuscript, *Clinical Viability of Magnetic Bead Implants in Muscle* (Taylor et al., 2022), we address the salient aspects of the clinical viability of magnetomicrometry. Namely, we describe the manufacturing process for the implants and implant insertion devices, assessed gait metrics and bead migration, and verified the biocompatibility of the implants via intramuscular implantation, cytotoxicity, sensitization, and intracutaneous irritation testing. This work provides further details on the biocompatibility of the magnetic bead implants and insertion devices described in that manuscript.

In this work, we verify the non-pyrogenicity of the magnetic bead implants and submit the implants to a full chemical characterization and toxicological risk assessment. Further, we validate the cleaning efficacy, steam sterilization, and dry time of the magnetic bead insertion device. We believe these results to be valuable to further scientific progress in the use of magnetic bead implants for human-machine interfacing.

## Materials and Methods

### Pyrogenicity Testing

To perform further biocompatibility testing on the device under good laboratory practice (GLP) compliance (USFDA, Code of Federal Regulations, Title 21, Part 58 -Good Laboratory Practice for Nonclinical Laboratory Studies), we submitted fully manufactured magnetic bead sets and insertion devices to WuXi AppTec for pyrogenicity testing. All magnetic beads used in the testing were deployed from the magnetic bead cartridges using the insertion device, and the testing was conducted in compliance with international standard ISO 10993-12:2012, Biological Evaluation of Medical Devices, Part 12: Sample Preparation and Reference Materials.

The pyrogenicity testing protocols were reviewed and approved by the WuXi AppTec IACUC prior to the initiation of testing. Eight rabbits were used for pyrogenicity testing in this work. Albino rabbits (Oryctolagus cuniculus, young adult, female) were obtained from Charles River Laboratories and maintained in the WuXi AppTec animal facility according to NIH and AAALAC guidelines on an ad libitum (except during the test period) water and certified commercial feed diet.

To test for the induction of a febrile response, magnetic beads were extracted at a ratio of 3 cm^2^ / 1 mL (surface area per volume) into each of 150.0 and 250.3 mL of 0.9% normal saline (1592 and 2656 magnetic beads for an initial test and a continued test, respectively). These extractions were freshly prepared for corresponding test phases and were performed over 72 hours at 50°C, with agitation during the extraction, then warmed to 37°C immediately before use.

In an initial test, 10 mL/kg of the extraction was slowly injected into the marginal ear vein in each of the first three rabbits. An automated temperature recorder measured a baseline rectal temperature no more than thirty minutes before the injection and the maximum temperature between one and three hours post-injection. This same testing was then performed in a continued test on the five additional rabbits. A temperature increase was calculated for each animal by subtracting the baseline temperature from the maximum temperature and rounding all negative temperature differences up to zero. If no temperature increase greater than 0.5°C had been observed in any of the animals in the initial test, the magnetic beads would have met the requirements of the test immediately following the initial test. The measurement of a temperature increase greater than 0.5°C in at least one animal of the initial test was considered grounds for performing a continued test, and a continued test was performed. As directed in the pyrogenicity testing procedure, fewer than three temperature increases greater than 0.5°C and a summed temperature increase of less than 3.3°C among the total of eight animals used in the initial and continued test were considered to indicate that the magnetic bead implants did not elicit a pyrogenic response. This pyrogenicity testing complied with United States Pharmacopeia (USP) Pyrogen Test Procedure, Section 151, with sample-specific preparation and extraction modifications as needed.

### Chemical Characterization and Toxicological Risk Assessment

To evaluate the toxicological potential of the magnetic bead implants as permanent implant devices, we submitted fully manufactured magnetic bead sets and insertion devices to Jordi Labs for chemical characterization. Gradient Corp then evaluated the chemical characterization results in a toxicological risk assessment. All magnetic beads used in the testing were deployed from the magnetic bead cartridges using the insertion device, and the testing was conducted in compliance with ISO 10993-12:2021, Biological Evaluation of Medical Devices, Part 12: Sample Preparation and Reference Materials. The chemical characterization was conducted in compliance with ISO 10993-18:2020, Biological evaluation of medical devices — Part 18: Chemical characterization of medical device materials within a risk management process.

For the performance of the chemical characterization, magnetic beads were extracted at a ratio of 3 cm^2^ / 1 mL (surface area per volume) into four 4.93 mL borosilicate vials each of purified water, ethanol, and hexane (12 sets of 48 magnetic beads – 576 beads total). Three of the vials for each extraction solvent were used to perform analysis in triplicate, while one vial was dedicated to gravimetric analysis. These extractions were performed by exhaustive submersion, with agitation during the extraction. The first extraction was performed over 72 hours at 50°C, while subsequent rounds would have been repeated over 24 hours at 50°C until the gravimetric analysis showed that non-volatile residue (NVR) was less than 10% the first extraction. However, all extractions met the exhaustive extraction criteria after the first round, with a measured total residue of less than 0.002 mg/device (less than the reporting limit of 0.1 mg), so the exhaustive extraction was stopped after one round. A control extraction was identically performed for each extraction solvent.

For the detection, identification, and quantitation of volatile and non-volatile organic compounds from the magnetic bead extractions, the extractions were analyzed by gas chromatography mass spectrometry (GCMS) and quadrupole time of flight liquid chromatography mass spectrometry with ultraviolet-visible and charged-aerosol detection (QTOF-LCMS-UV-CAD). The extractions were analyzed by head-space GCMS (HSGCMS) for volatile organic compounds. Elemental extractables were quantified using inductively coupled plasma mass spectrometry (ICP-MS). An analytical evaluation threshold (AET) was calculated as directed in the ISO standard, using a dose-based threshold of 1.5 ug/day and a maximum number of medical devices of sixteen. A dilution factor of four was used for QTOF-LCMS-UV-CAD, giving an AET concentration of 0.228 µg/mL, and a dilution factor of five was used for GCMS, giving an AET concentration of 0.182 µg/mL. A multidetector approach was used to reduce the effects of relative response factor variation (Jordi et al., 2020) and to provide an uncertainty factor to adjust the AET.

For the assessment of the identified chemicals for toxicological risk, an allowable limit for each chemical was calculated according to ISO Standard 10993-17:2002 using a minimum expected body weight of 58 kg. An exposure level was then conservatively calculated for each chemical using the assumption that the devices would daily release the maximum amount of chemical measured across all three solvents following the 72-hour exhaustive extraction and multiplying this by an expected maximum of sixteen devices per patient. The allowable limit was divided by this exposure level to give a margin of safety for each chemical. A margin of safety greater than 10 for compounds and greater than 1 for elements was considered to indicate low toxicological risk for each evaluated chemical for systemic toxicity, mutagenicity, carcinogenicity, and reproductive and developmental toxicity.

### Efficacy of Cleaning the Magnetic Bead Insertion Device

To test the efficacy of manually cleaning the magnetic bead insertion device for healthcare settings, we submitted three magnetic bead insertion devices to WuXi AppTec for cleaning efficacy testing.

We used the following instructions for use as the proposed cleaning process for the magnetic bead insertion devices: Disassemble the device into its four components, soak the components in a neutral-pH enzyme cleaner (Alconox Tergazyme, 1% solution at room temperature) for twenty minutes, and manually clean each component with scrub brushes and straw cleaners. Then, rinse each component with tap water. Next, sonicate the device components in a neutral-pH detergent (Alconox Luminox, 3% solution at room temperature) for ten minutes, and rinse each component of the device with tap water for one minute, then repeat the sonication and rinsing steps once more. Finally, wipe the devices with clean disposable non-shedding wipes (Kimwipes) and transfer the disassembled devices immediately to autoclave pouches for sterilization.

A full simulated-use cycle (See Figure A1) was defined as soiling each device, cleaning the devices with a worst-case cleaning process, then running the devices through a full autoclave cycle. To soil the devices, 2 mL of prepared artificial soil (ATS2015 soil – Healthmark) was spread on gloved hands, and the soiled gloved hands were used to handle and manipulate the device to spread the soil across all surfaces. The devices were then left to air-dry for a minimum of two hours. The worst-case cleaning process was chosen by decreasing the time of each step of the above proposed cleaning process (ten minutes of soaking in 0.5% enzyme cleaner, five-minute sonications in 1.5% solution detergent, and forty-five-second rinses), then allowing the devices to air-dry instead of wiping them down. Devices were autoclaved in a three-minute pre-vacuum cycle at 134°C, followed by a thirty-minute dry time.

**Figure A1:**
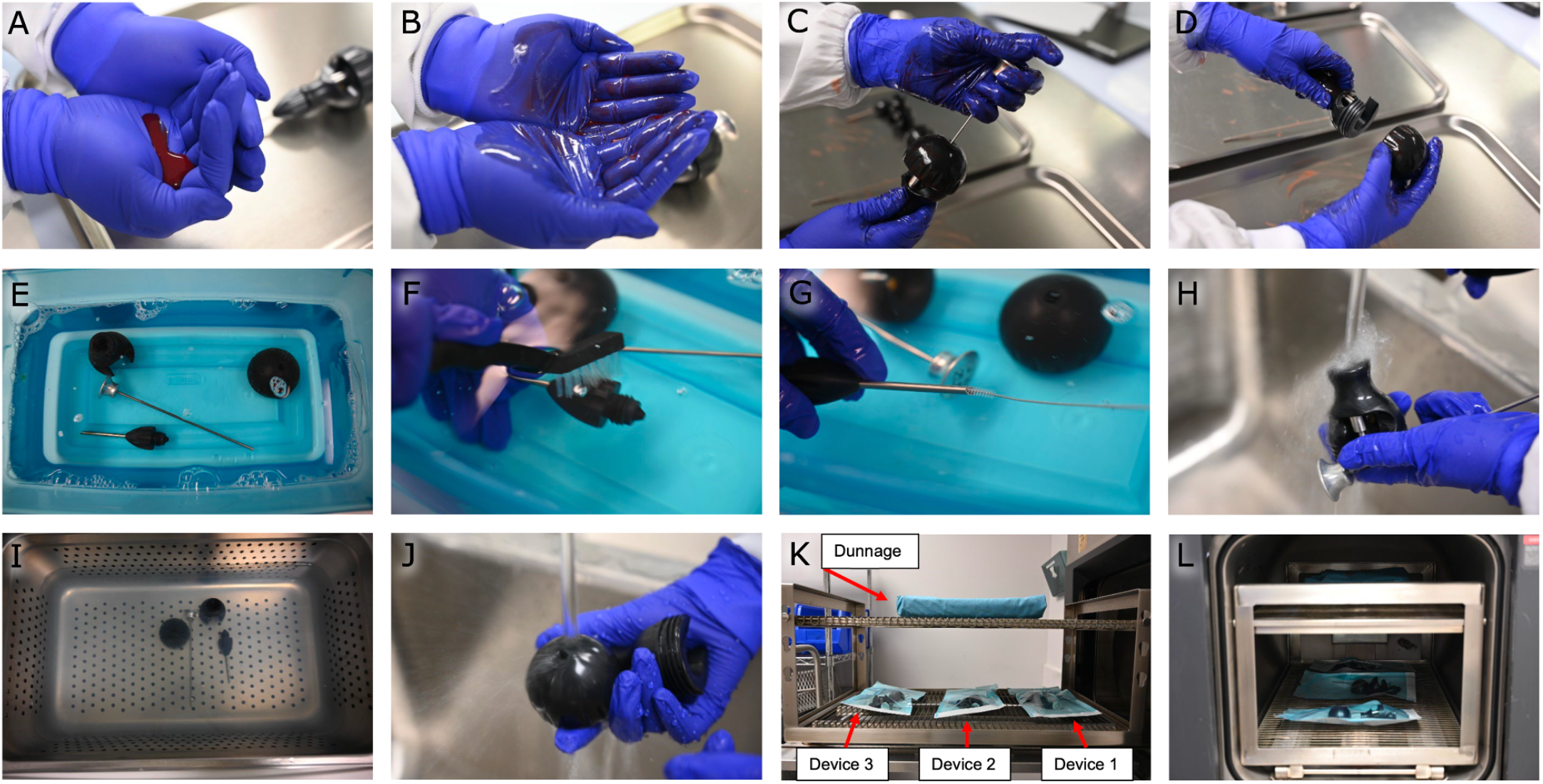
Cleaning Efficacy Study. (A) Preparation of artificial soil. (B) Spreading the artificial soil on gloved hands. (C, D) Handling and manipulating the device with soiled gloved hands. (E) Soaking the device in an enzyme cleaner. (F, G) Manually cleaning the device with scrub brushes and straw cleaners. (H) Rinsing the device with tap water. (I) Sonicating the device. (J) Rinsing the device with tap water. (K) Placing the devices on the autoclave rack with dunnage overhead. (L) Autoclaving the devices at 134°C. For the testing of remaining soil on the devices, each disassembled device was hand-shaken in an extraction bag with 200 mL of water for one minute, then sonicated in the bag for ten minutes, and hand-shaken for one additional minute. The sample extract was then tested for residual protein (via Micro BCA) and total organic carbon using quantitative colorimetric test methods.

One device served as a negative device control before any other testing occurred. The negative device control was cleaned according to the worst-case scenario (see Subfigures E-J of Figure A1) and then tested for remaining soil to verify that the measured soil quantities were at or slightly above negative sample controls. All three devices then served as positive device controls. The positive device controls underwent six full simulated-use cycles (see Subfigures A-L of Figure A1). They were then soiled once (see Subfigures A-D of Figure A1) and extracted three times to verify that a correction factor was unnecessary to account for recovery efficiency. Positive sample controls were then used to ensure the validity of the soil test assays.

For the testing of cleaning efficacy when manually cleaning the devices, two manual cleaning efficacy cycles were then performed on all three devices by soiling the devices, then cleaning them following the worst-case cleaning process (see Subfigures A-J of Figure A1). After each manual cleaning efficacy cycle, the devices were visually verified to be clean on all surfaces, then tested for remaining soil as described above. The cleaning efficacy study was considered to have met the acceptance criteria if the remaining soil per total device surface area was measured to be less than 6.4 μg/cm^2^ for protein and less than 12.0 μg/cm^2^ for total organic carbon.

### Steam Sterilization and Dry Time Validation for the Magnetic Bead Insertion Device

Steam sterilization and dry time were also validated by WuXi AppTec.

For the validation of the process for sterilizing the magnetic bead insertion device, the three magnetic bead insertion devices were inoculated with five strips each, with each strip containing 10^6^ *Geobacillus stearothermophilus* bioindicator spores (BI strip -Mesa Labs) having a decimal reduction value of two minutes at 121°C. As shown in subfigures A-E of Figure A2, the bioindicator strips were fed into the titanium shaft, the cap, through the center hole of the handle, and wrapped around the distal and proximal ends of the pushrod. The disassembled devices were then placed in autoclave pouches to be sterilized for a one-half cycle (1.5 minutes) at 134°C with dunnage overhead to simulate a full autoclave load. An additional bioindicator strip was handled similarly but was left outside the autoclave as a positive control. All bioindicator strips were retrieved and incubated for seven days at 55-60°C while fully immersed in Soybean-Casein Digest Broth. The steam sterilization validation was considered to indicate a 10^6^ sterility assurance level (SAL) for a full 134°C autoclave sterilization cycle if the sterility test results for the test samples and positive controls were all found to be negative and positive, respectively. This steam sterilization process validation was conducted in compliance with ISO 17665-1: 2006, Sterilization of health care products --Moist heat --Part 1: Requirements for the development, validation and routine control of a sterilization process for medical devices.

**Figure A2:**
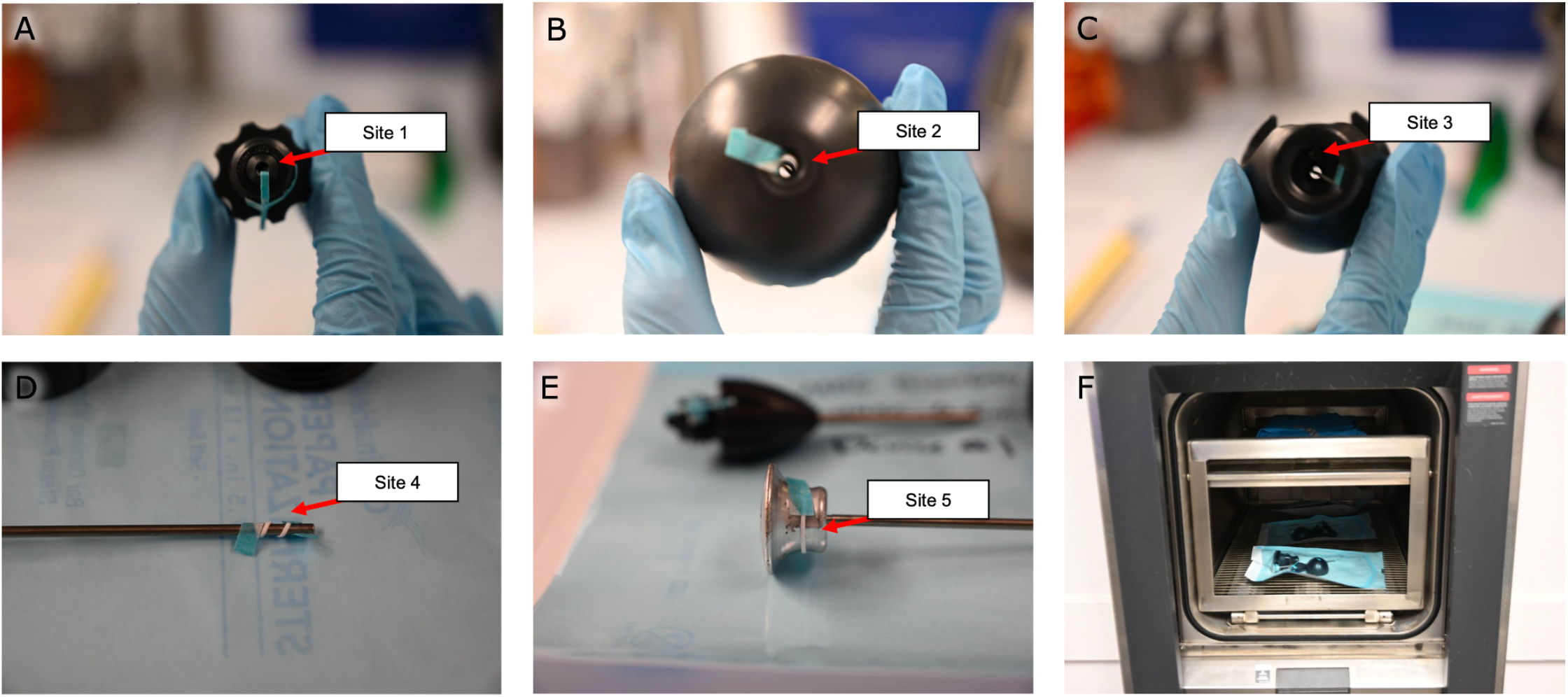
Steam Sterilization Study. Bioindicator strips were fed (A) into the titanium shaft, (B) the cap, (C) through the center hole of the handle, and wrapped around the (D) distal and (E) proximal ends of the pushrod. (F) The disassembled devices were autoclaved for a half-cycle before being tested for sterility. Note the dunnage on the top rack used to simulate a full autoclave load.

For validation of the post-sterilization dry time of the magnetic bead insertion device, the three magnetic bead insertion devices were individually weighed separate from their autoclave packaging, packaged, then placed in an autoclave with dunnage overhead as was performed in the sterilization validation to simulate a full autoclave load (as shown in subfigure F of Figure A2). The devices were autoclaved at 134°C for three minutes, then dried in the autoclave at a maintained temperature of 134°C for thirty minutes. Each device was unpackaged, and each device and packaging were inspected separately for residual moisture and weighed. This process was performed three times. The post-sterilization dry time study was considered to have met the acceptance criteria if no visible moisture was observed and neither the device nor the packaging weight was measured to have increased by more than 3%.

## Results

### Pyrogenicity Testing

The pyrogenicity test resulted in a passing score, indicating that the magnetic bead implants of this study are non-pyrogenic. In the initial test, a temperature increase of 0.5°C was measured in one animal, suggesting continued testing. In the continued test, no additional temperature increases greater than 0.5°C were observed, and the summed temperature increase was 1.1°C (see Table A1).

**Table A1:** Pyrogenicity Testing Results. The following table lists the baseline temperatures for each rabbit in the 30 minutes before injection of the magnetic bead extraction, the maximum temperatures recorded for each rabbit 1-3 hours post-injection, and the corresponding temperature increases (rounded up to zero), along with their individual pass/fail designations. The summed temperature increase across all eight rabbits is shown. No more than three animals received individual fail designations, and the summed temperature increase was less than 3.3°C, so the study result is non-pyrogenic.

|  | Rabbit | Baseline<br>Temperature (°C) | Maximum<br>Temperature (°C) | Temperature<br>Increase (°C) | Pass/Fail |
| --- | --- | --- | --- | --- | --- |
| Initial<br>Test | 1 | 39.2 | 39.2 | 0 | Pass |
|  | 2 | 39.0 | 39.2 | 0.2 | Pass |
|  | 3 | 39.0 | 39.5 | 0.5 | Fail |
| Continued<br>Test | 4 | 39.4 | 39.5 | 0.1 | Pass |
|  | 5 | 39.2 | 39.3 | 0.1 | Pass |
|  | 6 | 39.2 | 39.2 | 0 | Pass |
|  | 7 | 39.5 | 39.6 | 0.1 | Pass |
|  | 8 | 39.4 | 39.5 | 0.1 | Pass |
| Summed Temperature Increase (°C) and Study Result: |  |  |  | 1.1 | Pass (non-pyrogenic) |

### Chemical Characterization and Toxicological Risk Assessment

The chemical characterization detected, identified, and quantified 14 compounds and 7 elements (see Table A2). As listed in Table A2, all margins of safety for the identified compounds were greater than 10 and all margins of safety for identified elements were greater than 1, supporting a conclusion of tolerable risk of systemic toxicity, genotoxicity, carcinogenicity, and reproductive and developmental toxicity to patients.

**Table A2:**
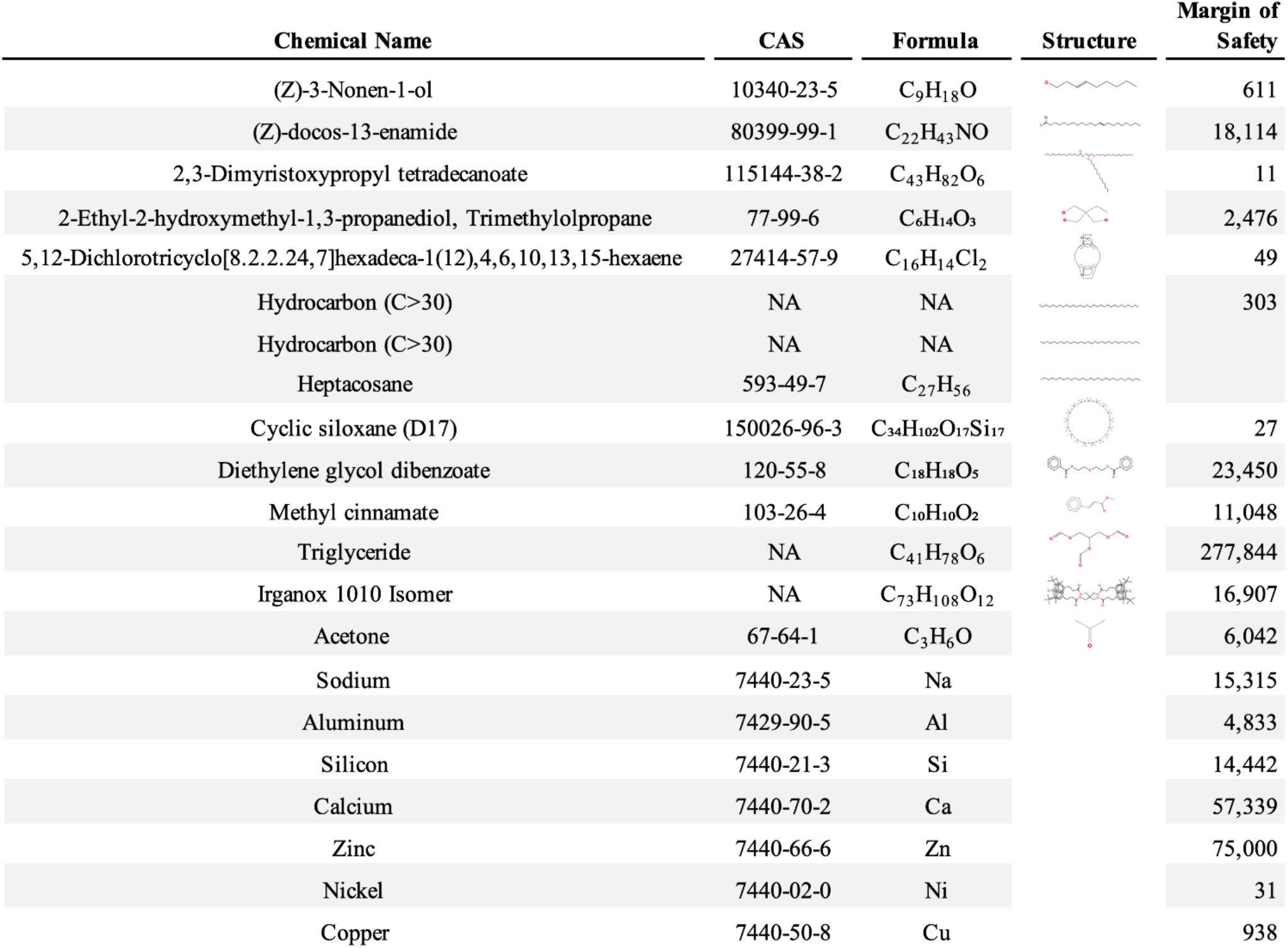
Chemical Characterization and Toxicological Risk Assessment Results. The table shows the chemicals found across all mass spectrometry analyses of all extractions of the magnetic beads, along with their CAS registry numbers, formulas, and structures. The three columns to the right of the table give the allowable limit for each chemical a patient can be safely exposed to, a conservatively calculated exposure level for that chemical, and a margin of safety ratio of the allowable limit and exposure level. All margins of safety are above 10 for compounds and above 1 for elements, supporting a conclusion of tolerable risk.

| Chemical Name | CAS | Formula | Structure | Margin of Safety |
| --- | --- | --- | --- | --- |
| (Z)-3-Nonen-1-ol | 10340-23-5 | C<sub>9</sub>H<sub>18</sub>O |  | 611 |
| (Z)-docos-13-enamide | 80399-99-1 | C<sub>22</sub>H<sub>43</sub>NO |  | 18,114 |
| 2,3-Dimyristoxypropyl tetradecanoate | 115144-38-2 | C<sub>43</sub>H<sub>82</sub>O<sub>6</sub> |  | 11 |
| 2-Ethyl-2-hydroxymethyl-1,3-propanediol, Trimethylolpropane | 77-99-6 | C<sub>6</sub>H<sub>14</sub>O<sub>3</sub> |  | 2,476 |
| 5,12-Dichlorotricyclo[8.2.2.24,7]hexadeca-1(12),4,6,10,13,15-hexaene | 27414-57-9 | C<sub>16</sub>H<sub>14</sub>Cl<sub>2</sub> |  | 49 |
| Hydrocarbon (C>30) | NA | NA |  | 303 |
| Hydrocarbon (C>30) | NA | NA |  |  |
| Heptacosane | 593-49-7 | C<sub>27</sub>H<sub>56</sub> |  |  |
| Cyclic siloxane (D17) | 150026-96-3 | C<sub>34</sub>H<sub>102</sub>O<sub>17</sub>Si<sub>17</sub> |  | 27 |
| Diethylene glycol dibenzoate | 120-55-8 | C<sub>18</sub>H<sub>18</sub>O<sub>5</sub> |  | 23,450 |
| Methyl cinnamate | 103-26-4 | C<sub>10</sub>H<sub>10</sub>O<sub>2</sub> |  | 11,048 |
| Triglyceride | NA | C<sub>41</sub>H<sub>78</sub>O<sub>6</sub> |  | 277,844 |
| Irganox 1010 Isomer | NA | C<sub>73</sub>H<sub>108</sub>O<sub>12</sub> |  | 16,907 |
| Acetone | 67-64-1 | C<sub>3</sub>H<sub>6</sub>O |  | 6,042 |
| Sodium | 7440-23-5 | Na |  | 15,315 |
| Aluminum | 7429-90-5 | Al |  | 4,833 |
| Silicon | 7440-21-3 | Si |  | 14,442 |
| Calcium | 7440-70-2 | Ca |  | 57,339 |
| Zinc | 7440-66-6 | Zn |  | 75,000 |
| Nickel | 7440-02-0 | Ni |  | 31 |
| Copper | 7440-50-8 | Cu |  | 938 |

### Efficacy of Cleaning the Magnetic Bead Insertion Device

The cleaning efficacy study met all acceptance criteria for the study (see Table A3), suggesting that the device can be safely cleaned in a healthcare setting according to the proposed cleaning process.

**Table A3:** Cleaning Efficacy Results. The soil per total device surface area was measured to be less than 6.4 μg/cm^2^ for protein and less than 12.0 μg/cm^2^ total organic carbon for all devices for both cleaning efficacy cycles, resulting in a passing score for the cleaning efficacy acceptance criteria.

| Device | 1 |  | 2 |  | 3 |  |  |
| --- | --- | --- | --- | --- | --- | --- | --- |
| Cleaning Efficacy Cycle | 1 | 2 | 1 | 2 | 1 | 2 |  |
| Protein (µg/cm<sup>2</sup>) | <0.6 | <0.6 | <0.6 | <0.6 | <0.6 | <0.6 | Pass |
| Total Organic Carbon (µg/cm<sup>2</sup>) | 1.1 | 1.1 | 1.2 | 1.2 | 1.9 | 1.5 | Pass |

### Steam Sterilization and Dry Time Validation for the Magnetic Bead Insertion Device

The steam sterilization validation met all acceptance criteria for the study, with all fifteen test samples testing negative and all three positive controls testing positive for the growth of *Geobacillus stearothermophilus*., indicating a 10^6^ sterility assurance level (SAL) for the magnetic bead insertion device after undergoing a full three-minute 134°C autoclave sterilization cycle.

The dry time validation met all acceptance criteria for the study. No moisture was observed on any of the devices during the dry time validation cycles. The weight of the devices and packaging decreased in all cases (see Table A4), indicating that the magnetic bead insertion device can be completely dried using a 30-minute autoclave dry time at 134°C following a full three-minute 134°C autoclave sterilization cycle.

**Table A4:**
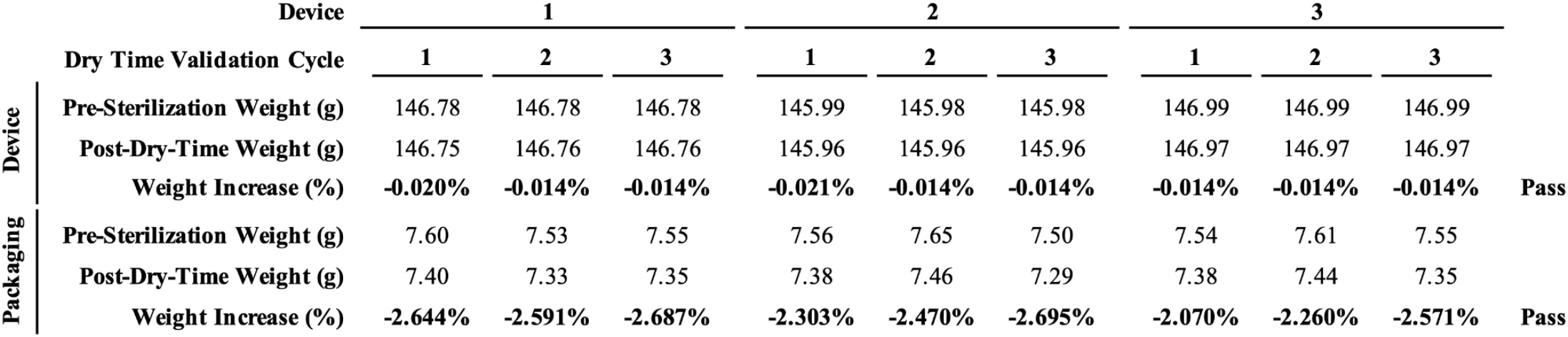
Dry Time Validation Results. The pre-sterilization and post-dry-time weights are shown for each device across all three dry-time validation cycles, along with the percent weight increase from before sterilization to after drying. The weight decreased in all cases, resulting in a passing score for the dry time validation.

## Discussion

This work further solidifies the clinical biocompatibility of the magnetic bead implants and their insertion devices, showing that the implants are non-pyrogenic and are considered to indicate low toxicological risk for each evaluated chemical for systemic toxicity, mutagenicity, carcinogenicity, and reproductive and developmental toxicity, and also that the insertion devices can be safely cleaned, sterilized, and dried in a healthcare setting. The results of this work, combined with the content of the main paper, will provide confidence for the safe use of these implants in an upcoming first-in-human clinical trial of magnetomicrometry.

## Acknowledgments

The authors thank, non-exhaustively, Zach Anderson, Gage Bateman, Steven Charlebois, Jenny Farr, Deborah Grayeski, Ted Heise, Mark Jordi, Kylie Kelley, Lindsey Reynolds, Hudson Santos, and Jessica Sohner for their helpful contributions, advice, suggestions, feedback, and support. Inclusion in this list of acknowledgments does not indicate endorsement of this work.

